# Re-evaluation of ethylene role in Arabidopsis cauline leaf abscission induced by water stress and rewatering

**DOI:** 10.1101/2021.09.27.462018

**Authors:** Shimon Meir, Sonia Philosoph-Hadas, Shoshana Salim, Adi Segev, Joseph Riov

## Abstract

Patharkar and Walker (2016) reported that cauline leaf abscission in Arabidopsis is induced by a cycle of water stress and rewatering, which is regulated by the complex of INFLORESCENCE DEFICIENT IN ABSCISSION (IDA), HAESA (HAE), and HAESA-LIKE2 (HSL2) kinases. However, they stated without presenting experimental results that ethylene is not involved in this process. Since this statement contradicts the well-established role of ethylene in organ abscission induced by a cycle of water stress and rewatering, our present study was aimed to re-evaluate the possible involvement of ethylene in this process. For this purpose, we examined the endogenous ethylene production during water stress and following rewatering, as well as the effects of exogenous ethylene and 1-methylcyclopropene (1-MCP), on cauline leaf abscission of Arabidopsis wild type. Additionally, we examined whether this stress induces cauline leaf abscission in ethylene-insensitive Arabidopsis mutants. The results of the present study demonstrated that ethylene production rates increased significantly in cauline leaves at 4 h after rewatering of stressed plants, and remained high for at least 24 h in plants water-stressed to 40 and 30% of system weight. Ethylene treatment applied to well-watered plants induced cauline leaf abscission, which was inhibited by 1-MCP. Cauline leaf abscission was also inhibited by 1-MCP applied during a cycle of water stress and rewatering. Finally, no abscission occurred in two ethylene-insensitive mutants, *ein2-1* and *ein2-5*, following a cycle of water stress and rewatering. Taken together, these results clearly indicate that ethylene is involved in Arabidopsis cauline leaf abscission induced by water stress.

**One sentence summary:** Unlike Patharker and Walker (2016), our results show that ethylene is involved in Arabidopsis cauline leaf abscission induced by water stress and rewatering, similar to leaf abscission in other plants.

## INTRODUCTION

Patharkar and Walker (2016) reported that Arabidopsis cauline leaves abscise as a result of water stress of 40% soil water content (SWC) followed by rewatering, and this process is regulated by the complex of INFLORESCENCE DEFICIENT IN ABSCISSION (IDA), HAESA (HAE), and HAESA-LIKE2 (HSL2) kinases. In this process, the IDA-HAE-HSL2 pathway operates through a mitogen-activated protein (MAP) kinase cascade MKK4/5 in a similar way as reported for Arabidopsis floral organ abscission after pollination (Jinn et al., 2000; Sung et al. 2008). Many of the genes required for Arabidopsis cauline leaf abscission triggered by a cycle water stress and rewatering were also required for abscission of Arabidopsis floral organs (Patharkar and Walker, 2016). However, in spite of the well-established role of ethylene in organ abscission (Jackson and Osborn, 1970; Taylor and Whitelaw 2001; Brown, 2006), these authors ruled out the possibility that ethylene triggers and/or is involved in cauline leaf abscission after a cycle of water stress and rewatering. This conclusion was expressed by their statement that “ethylene was not accumulated during cauline leaf abscission” (Patharkar and Walker, 2016). However, no information on the ethylene measurements was presented in this article, and no other means were used for ruling out the involvement of ethylene in this process, such as examination of ethylene-insensitive mutants.

Ethylene is an important candidate as a regulator of the process of leaf abscission induced by a cycle of water stress and rewatering. This is based on the massive literature showing that this process is ethylene-mediated in various plant species, including cotton (Jordon et al., 1972; Morgan et al., 1990), bean (Morgan et al., 1990), citrus (Tudela and Primo-Millo 1992; Agusti et al., 2012), and poplar (Street et al., 2006). Therefore, it is very surprising and even inconceivable that this well-established process, which was demonstrated in many plant species, will be differently regulated in the Arabidopsis model plant.

Distinct IDA family members from tomato and soybean (Tucker and Yang., 2012), citrus (Estornell et al., 2015), lychee (Ying et al., 2016), oil palm (Tranbarger et al., 2019), yellow lupine (Wilmowicz et al., 2018), and most recently from mango (Rai et al., 2021), were characterized and demonstrated to be involved in organ abscission. It should be noted that the expression of these *ida* gene family members increased in the above species by ethylene or ethephon application. This strengthens and establishes the suggestion that the IDA–HAE–HSL2 complex acts downstream of ethylene action, and that it is required for the later steps of organ execution rather than for the induction phase of the abscission process (Botton and Ruperti, 2019; Meir et al., 2019).

Furthermore, Patharkar and Walker (2016) and Patharkar et al. (2017) emphasized that they had never observed completely green leaves abscising after a drought treatment, and that the leaves always turned at least partially yellow before abscising. These data indicate that before the occurrence of abscission, the leaves were already in the process of senescence, in which ethylene plays an important role. Ethylene involvement as an inducer and/or accelerator of leaf senescence in a wide range of plant species, including Arabidopsis, has been established for at least 50 years of research (Grbić and Bleecker, 1995; Koyama 2014; Ueda et al., 2020).

In view of the above data, we believed that there is a need to examine this system more thoroughly and to re-evaluate the possible involvement of ethylene in the abscission process of Arabidopsis cauline leaves in response to a cycle of water stress and rewatering. For this purpose, we examined in this system of Arabidopsis WT plants the endogenous ethylene production, cauline leaf abscission in response to exogenous ethylene and/or to the ethylene action inhibitor, 1-methylcyclopropene (1-MCP). Cauline leaf abscission was also examined in two Arabidopsis ethylene-insensitive mutants.

The results of the present study clearly demonstrated that cauline leaf abscission induced by a cycle of water stress and rewatering was associated with elevated ethylene production in these leaves, and was inhibited by 1-MCP. Our results also showed that application of exogenous ethylene induced cauline leaf abscission even in unstressed plants. Furthermore, cauline leaf abscission was not detected in two ethylene-insensitive mutants following a cycle of water stress and rewatering. Based on these results and the available literature, it is concluded that ethylene is involved in Arabidopsis cauline leaf abscission induced by water stress and rewatering.

## RESULTS

A cycle of water stress to 40% of system weight and rewatering induced cauline leaf abscission, and about 30% of the premarked cauline leaves abscised after rewatering (Figure 1).

**Figure 1:**
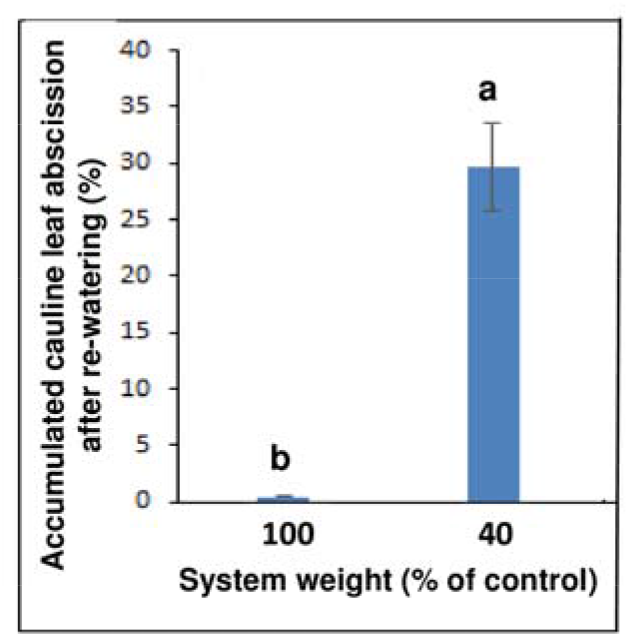
Effect of a cycle of water stress to 40% of system weight and rewatering on cauline leaf abscission. The system (plants + growing medium) was water-stressed until its weight reached 40% of the control (well-watered), and was rewatered thereafter. Cauline leaf abscission was monitored 2 and 5 days following rewatering, and the accumulated values are presented. Each bar represents means of three replicates ± SE, each including 10-12 plants. Different letters above bars denote statistical differences between treatments (*P* ≤ 0.01).

The possible involvement of ethylene in cauline leaf abscission in this system was first examined by measuring the ethylene production during various degrees of water stress and during rewatering. The data presented in Figure 2 show that during the water stress, the ethylene production rates did not differ significantly from the basic ethylene production rate of well-watered plants, which was about 1.8 nL g^-1^ h^-1^. On the other hand, 24 h after rewatering, when the abscission of the cauline leaves started, the ethylene production rates increased significantly by about 2.5-fold. We further examined the time course of ethylene production rates in cauline leaves during 24 h following rewatering. The data presented in Figure 3 show that 4 h following rewatering an increase in the ethylene production rates was recorded at all the water stress degrees. When the water stress was moderate, i.e. 60% (Figure 3A) or 50% (Figure 3B) of system weight, the increase in the ethylene production rates was transient, and at 6 h after rewatering the ethylene production rates returned to their basal level. On the other hand, when the water stress was stronger, i.e. 40% (Figure 3C) or 30% (Figure 3D) of system weight, the ethylene production rates continued to increase at 6 h, and were about 4.5- and 3.5- folds higher than the basic rates, respectively, and remained high 24 h following rewatering. These data further confirm the results presented in Figure 2 that the ethylene production rates of the cauline leaves increased only after rewatering. It should be noted that in these series of experiments, unlike the results presented in Figure 2, the ethylene production rate at 60% of system weight reached a relatively high level of 4.5 nL g^-1^ h^-1^ before rewatering (Figure 3A). This rate was about 2-fold higher than the ethylene production rates obtained at this time point at the other water stress degrees (Figures 3B, C, D).

**Figure 2:**
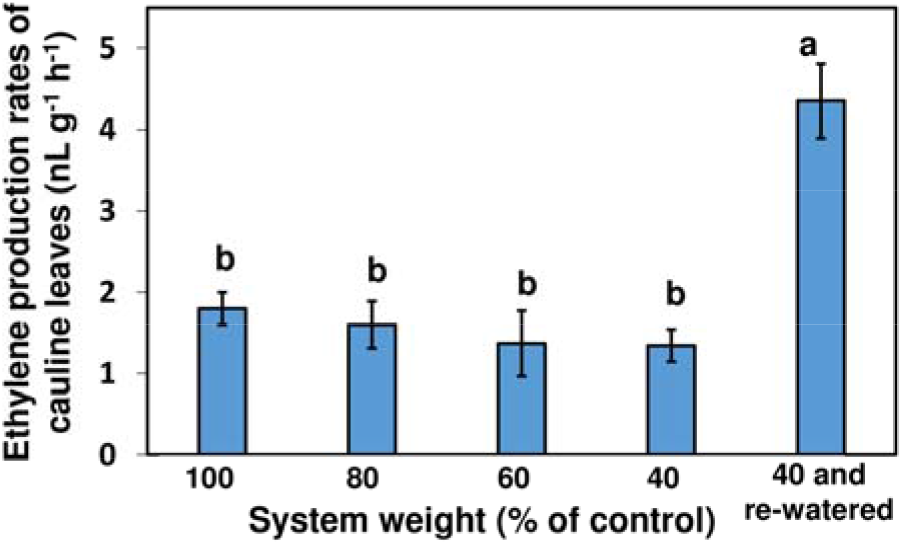
Effect of water stress to 80-40% of system weight and rewatering of the system stressed to 40% on ethylene production of cauline leaves. Ethylene production rates were monitored in cauline leaves detached from the plants at the indicated degrees of water stress, and 24 h following rewatering of the system stressed to 40%. Bars represent means of four replicates + SE, each including four detached leaves. Different letters above bars denote statistical differences between treatments (*P* ≤ 0.05).

**Figure 3:**
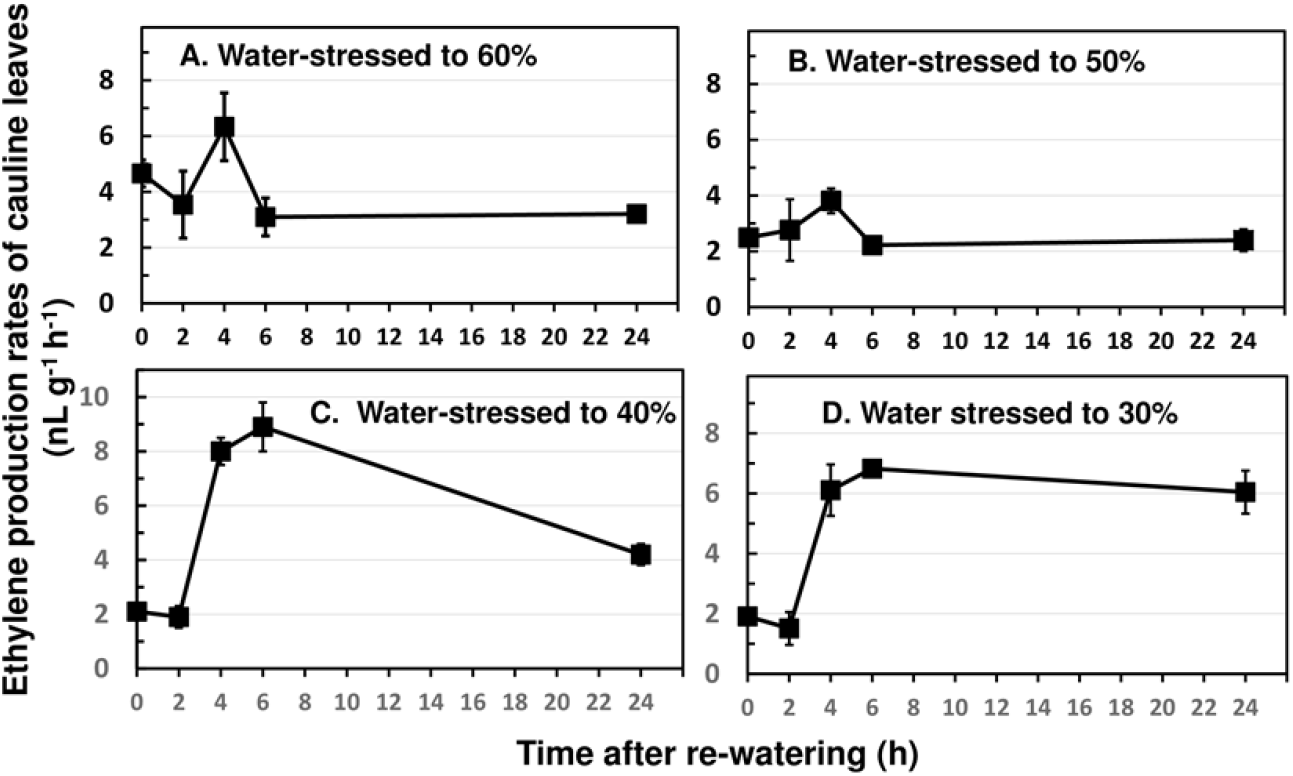
Time course of ethylene production in cauline leaves of plants water-stressed to 60% (A), 50%(B), 40% (C), or 30% (D) of system weight during 24 h following rewatering. Ethylene production rates were monitored in leaves detached from the plants at the different time points following rewatering. The results represent means of four replicates ± SE, each including four leaves. Different letters above symbols denote statistical differences between the time points (*P* ≤ 0.05).

To further study the involvement of ethylene, we examined whether exogenous ethylene can induce cauline leaf abscission in well-watered plants. Indeed, exposure of well-watered plants to 10 μL L^-1^ ethylene for 20 h induced cauline leaf abscission of 10%, which was significantly inhibited by the 1-MCP pretreatment (Figure 4). These results indicate that exposure to ethylene can cause abscission of cauline leaves even without water stress.

**Figure 4:**
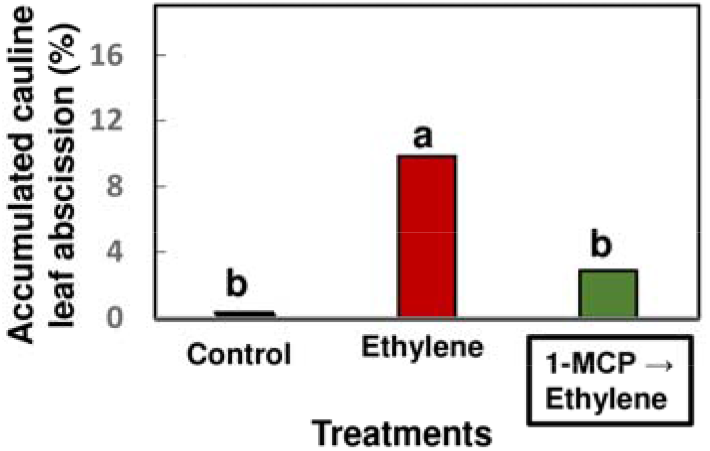
Effect of ethylene application with or without 1-MCP pretreatment to well-watered plants on cauline leaf abscission. Ethylene (10 μL L^-1^) was applied for 20 h to well-watered plants. 1-MCP (0.4 μL L^-1^) was applied for 4 h before the ethylene treatment. Cauline leaf abscission was monitored as detailed in Figure 1. Each bar represents means of three replicates, each including 10-12 plants. Different letters above bars denote statistical differences between treatments (*P* ≤ 0.05).

Exposure of the plants to 1-MCP at various time points during the application of water stress, either at 60% of system weight, or applied twice consecutively at 60% and then at 40% of system weight, reduced significantly the cauline leaf abscission after rewatering, as compared to control stressed plants without 1-MCP (Figure 5). When 1-MCP was applied twice consecutively at 60% and at 40% of system weight, an additive but not significant inhibitory effect was observed (Figure 5). We decided to apply 1-MCP at 60% of system weight because of the relatively high ethylene production rate observed in this treatment before rewatering (Figure 3A).

**Figure 5:**
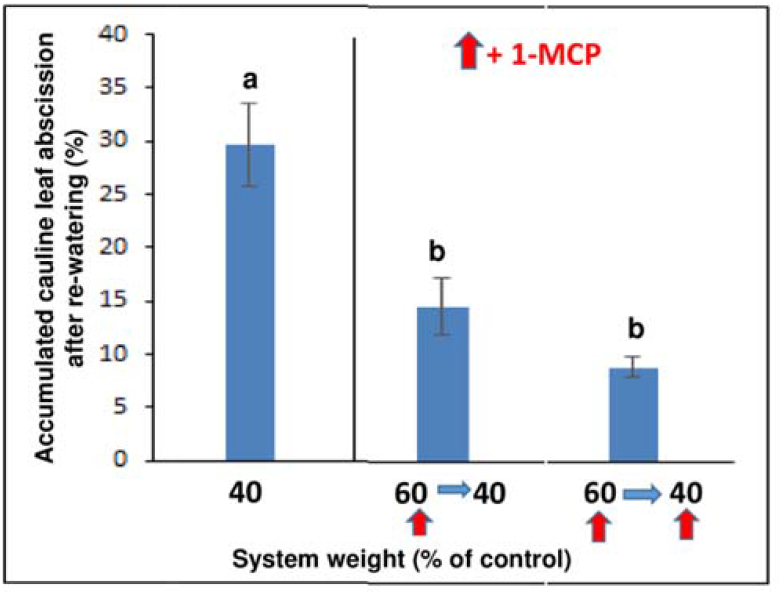
Effect of the timing of 1-MCP application at different water stress degrees before rewatering on cauline leaf abscission after rewatering. 1-MCP (0.4 μL L^-1^ for 4 h) was applied before rewatering at the system weights indicated by red arrows. The systems (plants + growing medium) were water-stressed until their weight reached 40% of the control (well-watered), and rewatered thereafter. Cauline leaf abscission was monitored as detailed in Figure 1. Each bar represents means of three replicates + SE, each including 10-12 plants. Different letters above bars denote statistical differences between treatments (*P* ≤ 0.05).

Finally, the role of ethylene as an inducer of Arabidopsis cauline leaf abscission was reinforced by examining the response of ethylene-insensitive mutants to a cycle of water stress and rewatering. The data presented in Figure 6 show that in the ethylene-insensitive mutants, *ein2-1* and *ein2-5*, no cauline leaf abscission occurred after a cycle of water stress to 40% of system weight and rewatering.

**Figure 6:**
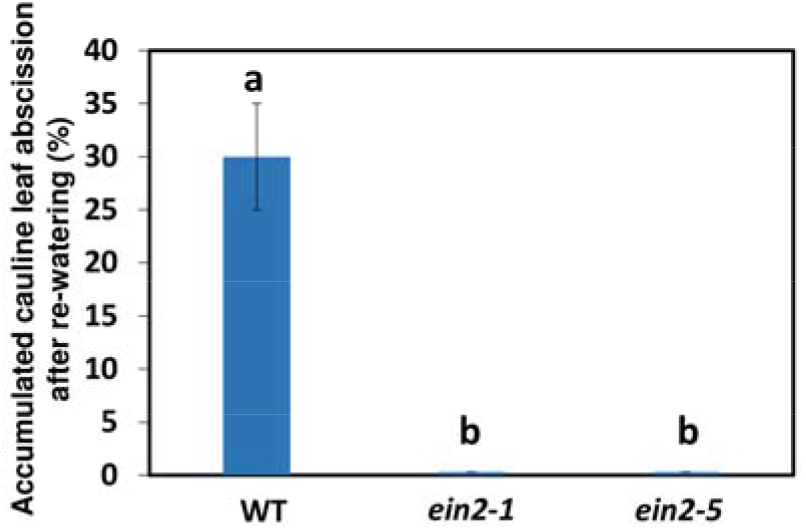
Effect of a cycle of water stress to 40% system weight and rewatering on cauline leaf abscission in Arabidopsis ethylene-insensitive mutants *ein2-1* and *ein2-5*. The systems (plants + growing medium) were water-stressed until their weight reached 40% of the control (well-watered), and were rewatered thereafter. Cauline leaf abscission was monitored as detailed in Figure 1. Each bar represents means of three replicates ± SE, each including 10-12 plants. Different letters above bars denote statistical differences between treatments (*P* ≤ 0.01).

Additionally, no yellowing of the cauline leaves could be detected in water-stressed plants of the ethylene-insensitive mutants (data not shown). These results further confirm that ethylene is necessary for exhibition of cauline leaf abscission and senescence induced by water stress.

## DISCUSSION

Ethylene is well known as a regulator of the process of leaf abscission induced by a cycle of water-stress and rewatering (Morgan et al., 1990; Tudela and Primo-Millo 1992; Street et al., 2006; Agusti et al., 2012). Hence, it is surprising that Patharkar and Walker (2016) ruled out the possible involvement of ethylene in this process in cauline leaves. The present study was aimed to examine their conclusion. In our experiments we used a water stress based on 40% of system weight, unlike a drought stress of 40% SWC used by Patharkar and Walker (2016). Nevertheless, we obtained similar results (Figure 1), showing a significant abscission of cauline leaves in response to 40% water stress and rewatering in our experimental system.

The results presented in Figures 2 and 3 clearly show that a cycle of water stress and rewatering increased the ethylene production of cauline leaves, which in turn induced their abscission. In a previous study on the effect of a cycle of drought stress and rewatering on citrus leaf abscission, it was shown that 1-aminocyclopropane (ACC) was synthesized in the roots during the drought period, and then transported to the leaves after rewatering, in which it was converted to ethylene, leading to leaf abscission (Tudela and Primo-Millo, 1992). The data presented in Figures 2 and 3 suggest, that also in the present study ACC was accumulated in the roots during the water stress in correlation with the degree of the stress, and *was* transported to the cauline leaves within 4 h after rewatering. Then, the transported ACC in the leaves was converted to ethylene, which induced the abscission process. Transport of ACC synthesized in the roots to upper organs was demonstrated in various plant systems. Bradford and Yang (1980) were the first to demonstrate that the transport of ACC from the roots to the shoots induced epinasty in tomato plants subjected to anaerobic root stress caused by flooding. Later, transport of ACC and its conjugated form malonyl-ACC from the roots to the shoots and leaves, was shown to occur during several abiotic stresses, including drought stress, as reviewed by Dubois et al. (2018).

Ethylene formed in the cauline leaves following water stress, could also induce the expression of the *ida* gene, and turn on the IDA-HAE-HSL2 pathway which is required for cauline leaf abscission (Patharkar and Walker, 2016). These authors reported that the *IDA* transcript increased in the AZ of the wilted plants at 40% SWC, and then it was further increased one day after rewatering (Patharkar and Walker, 2016). Ethylene was reported to induce *ida* expression in various plant species (Meir et al., 2019; Rai et al., 2021). The possibility that the increase of the *IDA* transcript in this system is ethylene-dependent has not yet been resolved. To get a clear answer to this question, the *IDA* transcript should be further analyzed in water-stressed plants of ethylene-insensitive mutants.

There are two main additional indications for the involvement of ethylene in cauline leaf abscission induced by water stress and rewatering: 1) inhibition of the process by the ethylene action inhibitor, 1-MCP (Figures 5); 2) no cauline leaf abscission occurred in the ethylene-insensitive mutants, *ein2-1 and ein2-5*, which underwent a cycle of water stress and rewatering (Figure 6). Thus, these data unequivocally indicate that ethylene is involved in the process of cauline leaf abscission induced by water stress and rewatering.

As reported by Patharkar and Walker (2016) and Patharkar et al. (2017), the cauline leaves always turned at least partially yellow before abscising. The same phenomenon was observed in the present study. This indicates that before abscission the leaves were already in the process of senescence, in which ethylene plays an important role (Grbić and Bleecker, 1995; Koyama, 2014). Indeed, the cauline leaves remained green in the water-stressed plants of the ethylene-insensitive mutants. It should be noted that the effect of the stress-induced endogenous ethylene production on cauline leaf abscission was much higher than the effect of exogenous ethylene applied to well-watered plants. Thus, the ethylene produced in water-stressed plants induced a 30% abscission of cauline leaves (Figures 1 and 5), whereas exogenous ethylene applied to well-watered plants induced only a 10% abscission (Figure 4). This difference might result from the fact that the stress-induced ethylene was produced after a cycle of water stress and rewatering applied to plants with cauline leaves that were in a yellowing process, while exogenous ethylene was applied to well-watered plants, in which the cauline leaves were still green. This suggests that the AZs of leaves that undergo the senescence process are more sensitive to ethylene than the AZs of green leaves. Nevertheless, 1-MCP pretreatment followed by ethylene application prevented the cauline leaf abscission in the well-watered plants, and this result indicates that the 10% abscission was induced by ethylene application.

## SUMMARY

In contrast to the report of Patharker and Walker (2016), our results indicate that ethylene is involved in Arabidopsis cauline leaf abscission induced by a cycle of water stress and rewatering. This conclusion is based on the following results:

1. Ethylene production rates significantly increased in cauline leaves 4 h after rewatering of 40%- or 30%-stressed plants, and remained high for at least 24 h.
2. Ethylene treatment applied to well-watered plants induced cauline leaf abscission.
3. The ethylene action inhibitor, 1-MCP, applied during the water stress (before rewatering) inhibited cauline leaf abscission.
4. No cauline leaf abscission occurred in two ethylene-insensitive mutants, *ein2-1* and *ein2-5*, following a cycle of water stress and rewatering.

## MATERIALS AND METHODS

### Plant Material, Growth Conditions and Treatments

The experiments were performed with *Arabidopsis thaliana* Columbia (Col-0) WT plants. In one experiment, the ethylene-insensitive mutants *ein2-1* and *ein2-5* were used. Following two days of vernalization at 4 °C, seeds were germinated on a half-strength Murashige and Skoog (MS) medium (Murashige and Skoog, 1962). Seven- to 10-day-old seedlings were transferred to 7×7×8 cm containers, containing 71.5 g of growing medium Green 7611 (Green Ltd., http://www.evenari.co.il), and grown at 22 °C under a 16 h/8 h light/dark cycle without fertilizer. During the growing period (about 5 weeks), the plants were irrigated every two days with tap water by flooding them at a height of 1 cm above the container base. The experiment was started with irrigation until saturation of the growing medium, carried out by complete coverage of the containers with water, followed by draining the excess water for half an hour and absorption of excess water at the base of the containers. At this stage, the systems (plants + growing medium) were weighed, and the weight was defined as 100% system weight at time zero. To induce water stress, irrigation was stopped, and the systems were weighed daily until their weight reached 80%, 60%, 50%, 40%, or 30% of their time zero weight. For rewatering the systems were irrigated to saturation as described above.

For application of ethylene, plants at various degrees of water stress were exposed after rewatering to 10 μL L^-1^ ethylene for 20 h in a 200-L closed chamber at 20 °C. For application of 1-MCP, plants were exposed before rewatering to 0.4 μL L^-1^ 1-MCP (Rimi Ltd. Sachets, Petah Tikva, Israel) for 4 h under the above conditions. In one experiment, the 1-MCP-treated plants were subsequently exposed to ethylene after rewatering as detailed above.

### Monitoring Abscission of Cauline Leaves

At the beginning of the experiment, healthy three cauline leaves were premarked on each inflorescence of the plant samples. Each premarked cauline leaf was touched gently, and leaves that fell off were counted as abscised. Cauline leaf abscission was monitored 2 and 5 days after rewatering, and the accumulated values of abscised leaves are presented. Cauline leaf abscission was expressed as the percentage of abscised leaves out of the total premarked cauline leaves on the plant samples at the beginning of the experiment. Experiments were performed with 3-4 replicates, each including 10-12 plants.

### Determination of Ethylene Production by Cauline Leaves

Samples of four cauline leaves were detached at different time points, weighted, and immediately enclosed in a sealed 5-mL syringe for 2 h at 20 °C. For determination of ethylene production, a 1-mL air sample was withdrawn from the syringe and injected into a gas chromatograph (Varian, Avondale, PA, USA), equipped with an alumina column and a flame ionization detector. Experiments were conducted with 4 replicates at each time point of each treatment.

### Statistical Analysis

Analysis of variance (ANOVA) (JMP Statistical Software; SAS Institute Inc., Cary, NC) was used to determine differences among treatments. Tukey’s post hoc was applied for mean comparisons of the significantly different treatments.

## List of author contributions

S.M. conceived the project and wrote the article with contributions of all the authors; S.M. and S.P.H. supervised the experiments, analyzed the data and wrote the draft version; S.S. and A.S. performed the experiments; J.R. supervised and completed the writing.

## Acknowledgements

Contribution from the ARO, Volcani Institute, Rishon LeZiyon, Israel. This research was funded by the Chief Scientist of the Israeli Ministry of Agriculture Fund, grant number 203-0898-11.

